# A rainfall-manipulation experiment with 517 *Arabidopsis thaliana* accessions

**DOI:** 10.1101/186767

**Authors:** Moises Exposito-Alonso, Rocío Gámez Rodríguez, Cristina Barragán, Giovanna Capovilla, Eunyoung Chae, Jane Devos, Ezgi S. Dogan, Claudia Friedemann, Caspar Gross, Patricia Lang, Derek Lundberg, Vera Middendorf, Jorge Kageyama, Talia Karasov, Sonja Kersten, Sebastian Petersen, Leily Rabbani, Julian Regalado, Lukas Reinelt, Beth Rowan, Danelle K. Seymour, Efthymia Symeonidi, Rebecca Schwab, Diep Thi Ngoc Tran, Kavita Venkataramani, Anna-Lena Van de Weyer, François Vasseur, George Wang, Ronja Wedegärtner, Frank Weiss, Rui Wu, Wanyan Xi, Maricris Zaidem, Wangsheng Zhu, Fernando García-Arenal, Hernán A. Burbano, Oliver Bossdorf, Detlef Weigel

**Affiliations:** Department of Molecular Biology, Max Planck Institute for Developmental Biology, Tübingen, Germany; Center for Plant Biotechnology and Genomics, Technical University of Madrid, Pozuelo de Alarcón, Spain; Institute of Ecology and Evolution, University of Tübingen, Tübingen, Germany

**Keywords:** *Arabidopsis thaliana*, Climate Change, Field experiment, Local adaptation

## Abstract

The gold standard for studying natural selection and adaptation in the wild is to quantify lifetime fitness of individuals from natural populations that have been grown together in a common garden, or that have been reciprocally transplanted. By combining fitness values with species traits and genome sequences, one can infer selection coefficients at the genetic level. Here we present a rainfall-manipulation experiment with 517 whole-genome sequenced natural accessions of the plant *Arabidopsis thaliana* spanning the global distribution of the species. The experiments were conducted in two field stations in contrasting climates, in the Mediterranean and in Central Europe, where we built rainout shelters and simulated high and low rainfall. Using custom image analysis we quantified fitness- and phenology-related traits for 23,154 pots, which contained about 14,500 plants growing independently, and over 310,000 plants growing in small populations (max. 30 plants). This large field experiment dataset, which associates fitness and ecologically-relevant traits with genomes, will provide an important resource to test eco-evolutionary genetic theories and to understand the potential evolutionary impacts of future climates on an important plant model species.

## Background and Summary

Darwin’s theory of evolution by natural selection states that when individuals of a population have distinct traits that improve their ability to survive and reproduce, and these are heritable, the population will change and adapt over generations^1^. This was formally described by R. A. Fisher^2^, stating that the higher the genetic variance in fitness, the higher the rate of adaptation of a species. Natural selectìon over morphological, physiological or other traits has been studied in a wide range of organisms^3–9^ using observational and experimental fitness measurements of multiple individuals in field conditions. However, studies that combine such measurements with knowledge on genome-wide variation are, in comparison, very rare^7–9^. This is surprising, given that they can enable the translation of selection coefficients to the genetic level and thus ultimately help us to understand whether traits will evolve over generations.

With climate change, the study of adaptation to the environment has acquired new importance. Predictions of climate change indicate not only that temperature will rise, but that also precipitation regimes will be altered, leading to more frequent and extreme droughts^10^, posing the critical question of whether populations will be able to adapt or will become extinct^11^. Field experiments where climate variables such as rainfall are manipulated can be used to address this question^12^.

Here we present a high-throughput field experiment with 517 whole-genome sequenced natural lines of *Arabidopsis thaliana*^13^ grown under rainfall-manipulation conditions at two field stations with contrasting climate. This experiment was designed to be of a sufficiently large scale to enable powerful genome-wide association analyses^14^ and to maximize the replicability of species-wide patterns, which increases with the diversity of genotypes included in an experiment^15^. This dataset will be invaluable for the study of natural selection and adaptation in the context of global climate change at the genetic level^16^, building on the genetic catalog of the 1001 Genomes Project^13^ and complementing the already published extensive set of traits measured in controlled growth chamber or greenhouse conditions^17,18^.

## Methods

### Accessions from the 1001 Genomes Project

The 1001 Genomes (1001G) Project^13^ has provided information on 1,135 natural lines or accessions and 11,769,920 SNPs and small indels called after re-sequencing (Fig. 1). To select the most genetically and geographically informative 1001G lines, we applied several filters: (1) First we removed the accessions with the lowest genome quality. We discarded those with < 10X genome coverage of Illumina sequencing reads and < 90% congruence of SNPs called from MPI and GMI pipelines^13^. (2) We removed near-identical individuals. Using Plink^19^ we computed identity by state across the 1,135 accessions. For pairs of accessions with < 0.01 differences per SNP (<100,000 variants approx.), we randomly selected one accession to include in our study. (3) Finally, we reduced geographic sampling ascertainment bias, as the sampling for 1001G was performed in neither a random nor a regularly structured scheme. Some laboratories provided several lines per location whereas others provided lines that were collected at least several hundred kilometres apart. Using each accession’s collection location, we computed Euclidean distances across the 1,135 accessions and identified all pairs that were apart less than 0.0001 Euclidean distance in degrees latitude and longitude (≪ 100 meters). From such pairs, we randomly selected one accession to remain. After applying criteria (1), (2), and (3), we obtained a final set of 523 accessions (Datasets 1 and 2). To bulk seeds for our rainfall-manipulati’on experiment and control for maternal effects, we first propagated accessions in controlled conditions. We stratified the seeds one week at 4°C, we sowed them in trays with industrial soil (CL-P, Einheitserde Werkverband e. V., Sinntal-Altengronau Germany) and placed them in a growth room with 16 h light and 23°C for one week. Trays were vernalized for 60 days at 4°C and 8h daylength. After vernalization, trays were moved back to 16 h light and 23°C for final growth and reproduction. This generated sufficient seeds for 517 accessions, which were later grown in the field in two locations (Fig. 1). Seeds originating from the same parents can be ordered from the 1001G seed stock at the Arabidopsis Biological Resource Center (CS78942).

**Figure 1.**
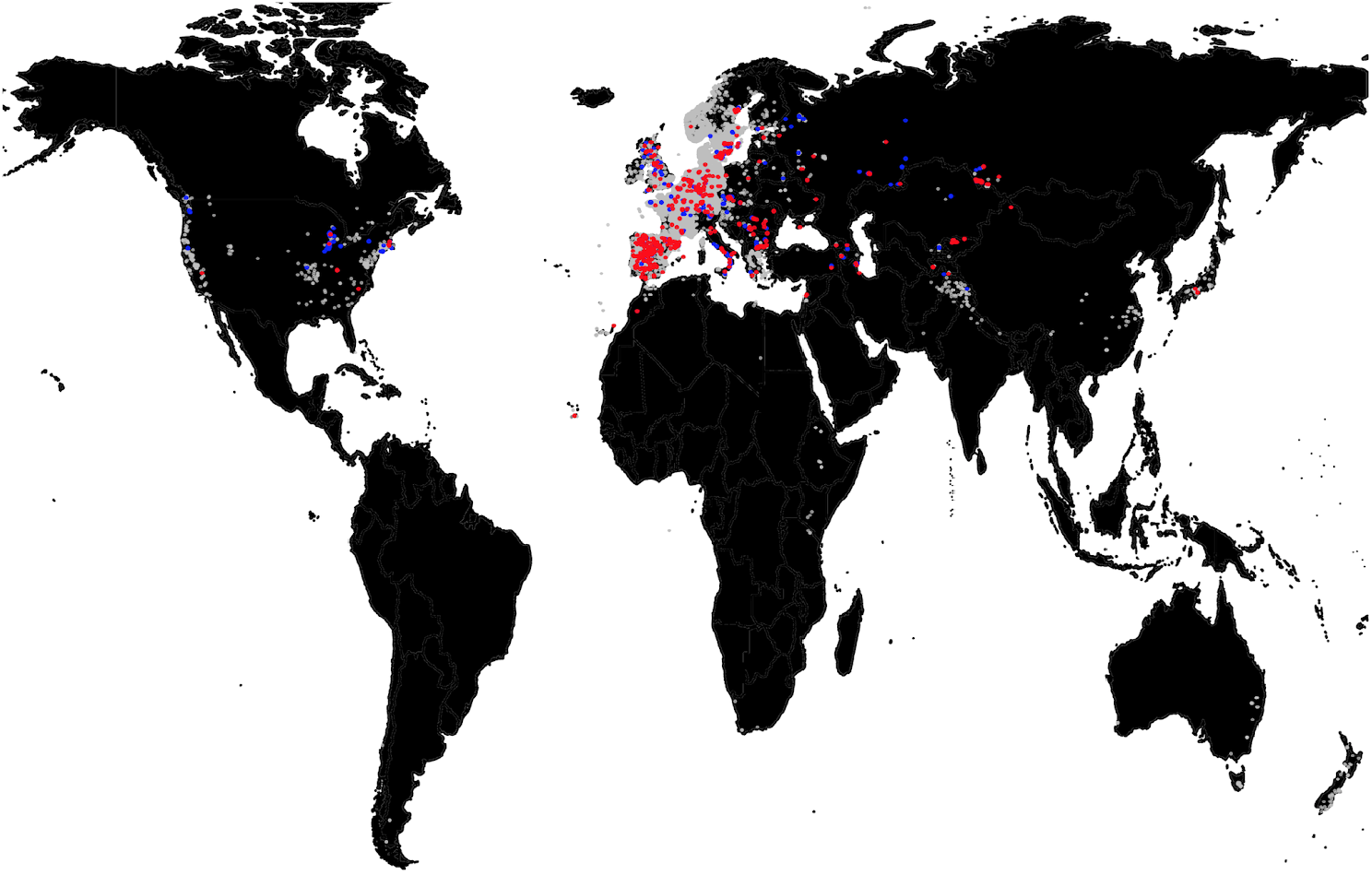
Geographic distribution of accessions. Locations of *Arabidopsis thaliana* accessions used in this experiment (red), 1001G accessions (blue), and all sightings of the species in gbif.org (grey).

### Field experiment design

#### Rainout shelter design

We built two 30 m × 6 m tunnels of PVC plastic foil to fully exclude rainfall in Madrid (Spain, <u>40.40805°N −3.83535°E</u>) and in Tübingen (Germany, <u>48.545809°N 9.042449°E</u>) (Fig. 2). The foil tunnels are different from a regular greenhouse in that they are completely open on two sides. Thus, ambient temperatures vary virtually as much as outside the foil shelter (see Environmental sensors section). In each location, we supplied artificial watering in two contrasting regimes: abundant watering and reduced watering. Inside each tunnel, we created a 4% slope, and four flooding tables (1 m × 25 m, Hellmuth Bahrs GmbH & Co KG, Brüggen, Germany) covered with soaking mats (4 l/m^2^, Gärtnereinkauf Münchingen GmbH, Münchingen, Germany) were placed on the ground in parallel to the slope. Water was able to drain at the lower end of the flooding table (Fig. 2). A watering gun was used to manually simulate rainfall from the top.

**Figure 2.**
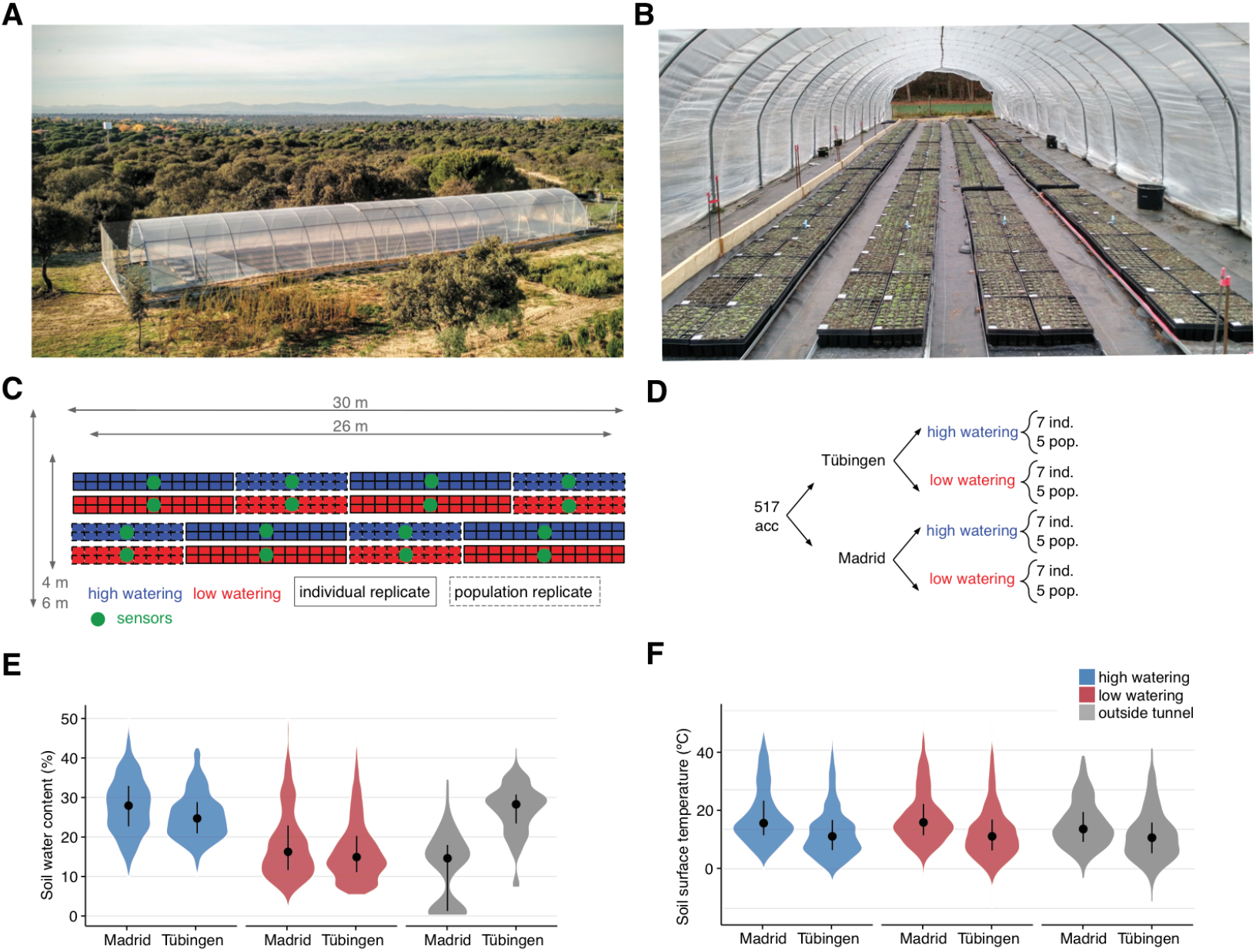
Field experiment design. (A) Aerial view of foil tunnel settings in Madrid and (B) view inside the foil tunnel in Tübingen. (C) Spatial distribution of blocks and replicates and (D) experimental design. (E) Soil water content and (F) soil surface temperature from the 34 sensors monitoring each experimental block and conditions outside the tunnel.

For sowing, we used potting trays with 8×5 cells (5.5 cm × 5.5 cm × 10 cm size) and industrial soil (CL-P, Einheitserde Werkverband e.V., Sinntal-Altengronau Germany). One genotype was sown per cell, excluding corner cells, to avoid extreme edge effects. We grew a total of 12 replicates per genotype per treatment: Five replicates were sown at high density, with 30 seeds per cell and without further intervention (“population replicate”). The remaining seven replicates were planted at low density (ca. 10 seeds) and one seedling was selected at random after germination (“individual replicate”). Excess individuals were culled. While the population replicates should more faithfully reflect survival from seed to reproduction, the individual replicates were useful to more accurately monitor flowering time and seed set.

We used a randomized incomplete block design (Fig. 2). One block of the 517 genotypes spanned 14.36 trays (36 cells/tray), and genotypes were randomized within each block. The design was identical in Madrid and Tübingen (Fig. 2).

#### Environmental sensors

Environmental variables — air temperature, photosynthetically active radiation (PAR) and soil water content — were monitored every 15 minutes for the entire duration of the experiment using multi-purpose sensors (Flower Power, Parrot SA, Paris, France). This enabled us to adjust watering depending on the degree of local evapotranspirati’on during the course the experiment. The sensors outside of the tunnel in Madrid (i.e. only natural rainfall) showed an interquartile range between 1% and 17% soil water content. This overlapped with the range of 10 to 22% water content of the drought treatment that we artificially imposed inside the tunnels in Madrid and Tübingen. The lower range of measurements in Madrid (outside sensor) is due to a lack of natural rainfall during the first two months of the experiment. In contrast, the sensor outside the tunnel in Tübingen recorded an interquartìle range of soil water content percentage of 22 to 27%, which was comparable to the high watering treatments in Tübingen and Madrid (from 20 to 33%). These values confirmed that our low and high watering treatment were not only different, but also that they mimicked natural soil water content at the two contrasting locatìons. Mean daily air temperatures (measured by the sensors at 5-10 cm above the soil surface every 15 minutes) were overall higher in Madrid (8-10°C) than in Tübingen (5-6°C), and the difference in temperature between the sensors inside and outside the tunnels was in both locatìons on average only 1°C (Table 1). The photosynthetìcally actìve radiation (PAR, wavelengths from 400 to 700 nm) had a median of 0.1 mol m^-2^ day^-1^ at night for all experiments. At mid-day (11:00-13.00 hrs), the median PAR in Madrid was 57.8 mol m^-2^ day^-1^ outside, and 45.7 mol m^-2^ day^-1^ inside the tunnel. In Tübingen, the median values were 29.0 outside, and 30.9 mol m^-2^ day^-1^ inside the tunnel.

**Table 1.** Summaries of environmental sensor measurements. A total of 34 sensors were placed in the different treatment blocks (low/high) as well as outside (out) of the foil tunnels (Fig. 2). The median (interquartìle) values of all sensors per treatment and location are shown.

| Site | Rainfall | Soil water content (%) | Air temperature (°C) |
| --- | --- | --- | --- |
| Madrid | out | 14.5 (1.09, 17.46) | 8.5 (5.34, 12.39) |
| Madrid | low | 16.1 (11.38, 22.51) | 10.0 (6.95, 15.13) |
| Tuebingen | low | 14.7 (10.76, 20.09) | 6.6 (3.27, 10.78) |
| Tuebingen | out | 27.7 (22.82, 30.50) | 5.6 (2.44, 9.54) |
| Tuebingen | high | 24.6 (20.73, 29.02) | 6.6 (3.27, 10.78) |
| Madrid | high | 27.8 (22.62, 33.00) | 9.8 (6.82, 15.13) |

#### Sowing and quality control

During sowing, contamination of neighboring pots with adjacent genotypes can occur for multiple reasons. In order to avoid such contamination, we chose a day with no wind and sowed seeds at 1-2 cm height from the soil. Additionally, we took care during the first days to be particularly gentle when using the watering gun to avoid seed-carryover (bottom watering by flooding was done regularly). We also tried to remove human error during sowing by preparing and randomizing 2 ml plastic tubes containing the seeds to be sown in the same layouts (5×8) as the destination trays. During sowing, each experimenter took a box at random and went to the corresponding labeled and arranged tray in the field (Fig. 2). This reduced the possibility of sowing errors. During vegetative growth, we could identify seedlings that resembled their neighbors or were located in the border between two pots and removed such plants as potential contaminants. We also used the homogeneity of flowering within a pot in the population replicates as a further indicator for contamination. When a plant had a completely different flowering timing or vegetative phenotypes did not coincide with the majority of plants in the pot, this plant was removed. After sowing and quality control, the total number of pots was 24,747 instead of the original 24,816 pots (99.7%).

### Field monitoring

#### Image analysis of vegetative rosettes

Top-view images were acquired every four to five days (median in both sites) with a Panasonic DMC-TZ61 digital camera and a customized closed dark box, the “Fotomatón” (Fig. 3), at a distance of 40 cm from each tray. In total, we imaged each tray at 20 timepoints throughout vegetative growth. The implemented segmentation was the same as in Exposito-Alonso et al.^20^, which relies on the Open CV Python library^21^. We began by transforming images from RGB to HSV channels. We applied a hard segmentation threshold of HSV values as (H=30-65, S=65-255, V=20-220). The threshold was defined after manually screening 10 different plants in order to capture the full spectrum of greens both of different accessions and of different developmental stages. This was followed by several iterations of morphology transformations based on erosion and dilation. For each of the resulting binary images we counted the number of green pixels.

**Figure 3.**
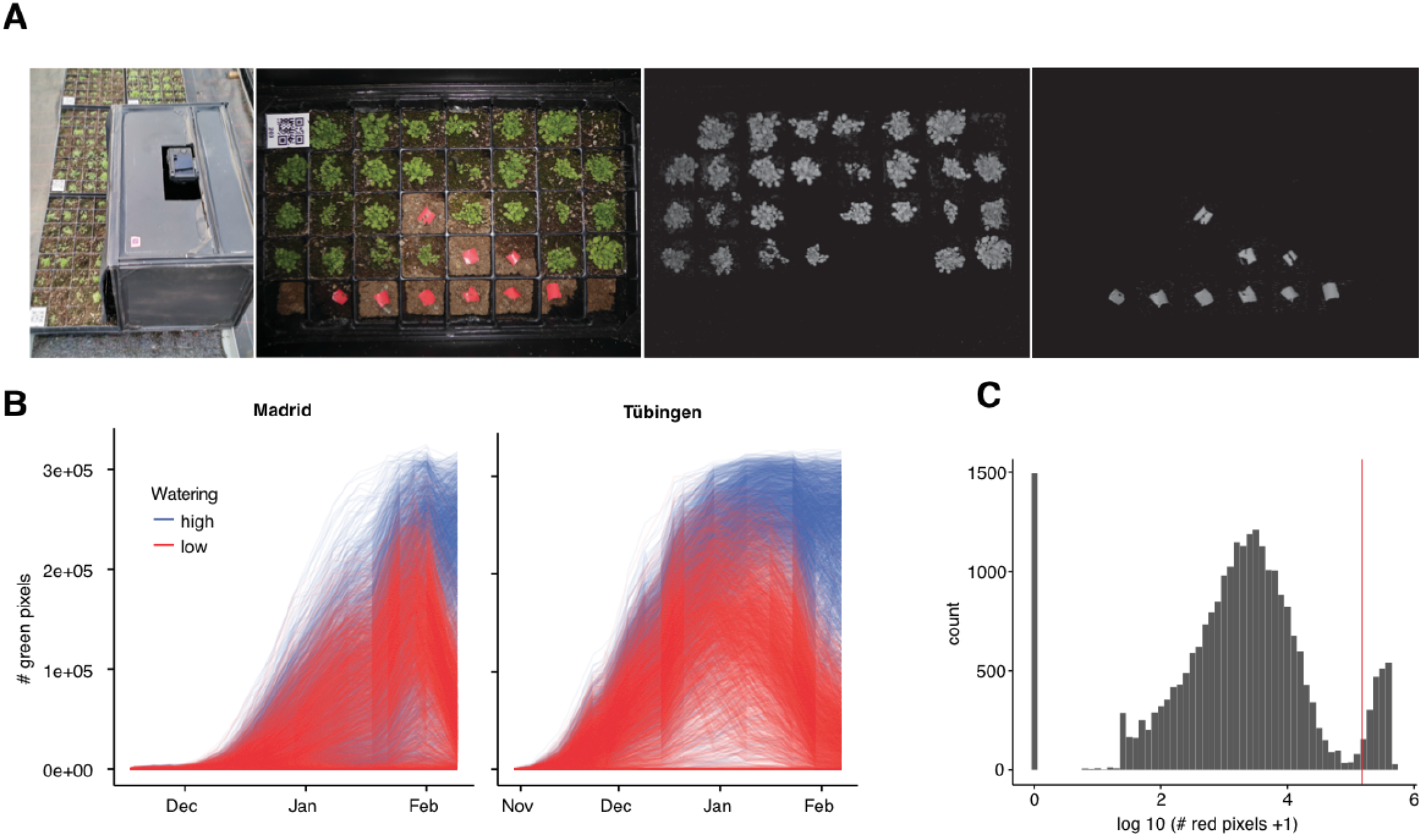
Rosette monitoring. (A) Customized dark box (“Fotomatón”) for image acquisition and example tray with the corresponding green and red segmentation. (B) Trajectories of number of green pixels per pot, indicating rosette area, for Madrid and Tübingen. (C) Distribution of the sum of red pixels per pot over all time frames. The red vertical line indicates the heuristically chosen threshold to define whether the pot actually had a red marker.

During field monitoring, we noticed that some pots were empty because seeds had not germinated. In these cases, we left a red marker in the corresponding pots, which could be detected in a similar way as the presence of green pixels (with threshold H=150-179, S=100-255, V=100-255). These pots were excluded from survival analysis as they did not contain any plants. An example of transformed images is shown in Fig. 3. The resulting raw data consist of green and red pixel counts per pot (Fig. 3). In order to detect the red markers automatically, we performed an analysis of variance between pots above and below a threshold of red pixels and finding the threshold that maximized this separation. This provided us with the threshold of red pixels above which a pot had a red marker (indicating an empty pot). As expected, the distribution of pixels was bimodal, making this identification straightforward (Fig. 3C).

We estimated germination timing by analysing trajectories (Fig. 3) of green pixels per pot, and identifying the first day that over 1,000 green pixels were observed in a pot (corresponding to a plant size of ~ 10 mm^2^, Fig. 4). The final dataset contained data for 22,779 pots — after the removal of pots with red labels — with a time series of green pixel counts.

**Figure 4.**
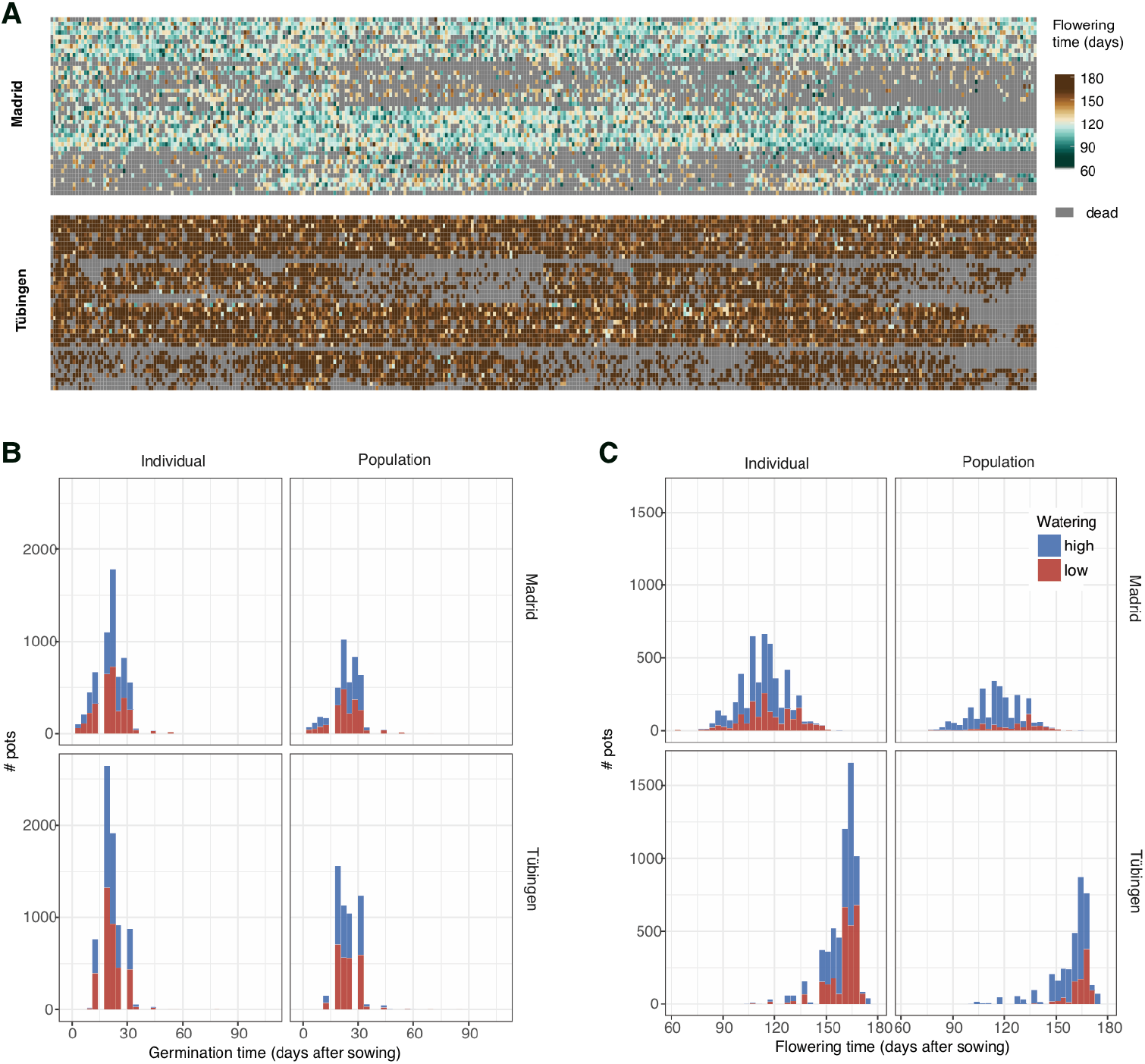
Flowering time distributions. (A) Flowering times per pot in the same spatial arrangement as in each tunnel (see Fig. 3). (B) Distribution of germination times. (C) Distribution of flowering times.

#### Manual recording of flowering time

We visited the experimental sites every 1-2 days and manually recorded the pots with flowering plants. Flowering time was measured as the day when the first white petals could be observed with the unaided eye. This criterion was chosen as sufficiently objective to reduce experimenter error. To keep track of previous visits and avoid errors, we labeled the pots where flowering had already been recorded with blue pins. To calculate flowering time, we counted the number of days from the date of sowing to the recorded flowering date (we did not use the inferred day of germination to avoid introducing modeling errors in the flowering time metric). Fig. 4 shows the raw flowering time data per pot in the original spatial distribution (Fig. 2) and the distribution of flowering time per treatment combination. Note that grey boxes in Fig. 4 are pots with plants that did not survive until flowering. In total, we gathered data for 16,858 pots with flowering plants.

#### Image analysis of reproductive plants

Once the first dry fruits were observed, we harvested them and took a final ‘studio photograph’ of the rosette and the inflorescence (Fig. 5). In total, we took 13,849 photographs. The camera sefflngs were the same as for the vegetative monitoring, but here we included an 18% grey card approximately in the same location for each picture in case *a posteriori* white balance adjustments would be needed. We first used a cycle of morphological transformations of erode-and-dilate to produce the segmented image (Fig. 5). This generated a segmented white/black image without white noise. Then, we used the thin (erode cycles) algorithm from the Mahotas Python library^22^ to generate a binary picture reduced to single-pixel paths — a process called skeletonisation (Fig. 5). Finally, to detect the branching points in the skeletonised image we used a hit-or-miss algorithm. We used customized structural elements to maximize the branch and end point detection (Fig. 5). This resulted in four variables per image: total segmented inflorescence area, total length of the skeleton path, number of branching points, and number of end points (Fig. 5).

**Figure 5.**
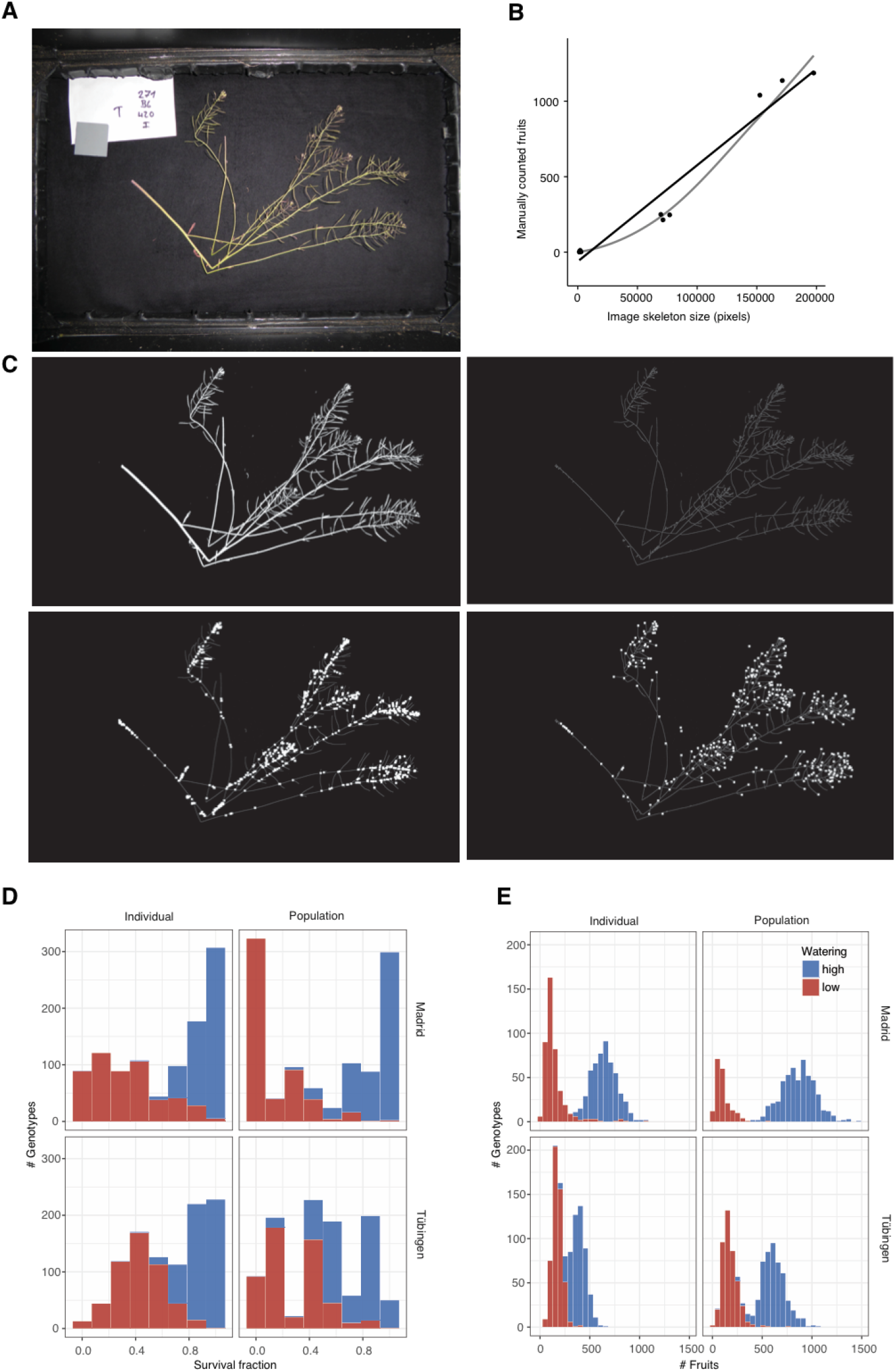
Inflorescence and seed set estimation. (A) Representative inflorescence picture. (B) Regression between the fruits of a few manually counted inflorescences and the inflorescence size calculated based on image processing. The four variables inferred in (C) accurately predicted the visually counted inflorescences as example (R^2^=0.97, n=11, P=10^−4^). (C) Resulting variables from image processing of (A): total segmented area (upper-left), skeletonized inflorescence (upper-right), branching points (lower-left), and endpoints (lower-right). Distribution of survival to reproduction (D) and fruits per plant (E) in the four environments.

#### Estimation of fruit and seed number

Although the study of natural selection is based on studying relative fitness, and total reproductive area might provide a good relative estimate, sometimes it is useful to have a proxy of the absolute fitness. In order to provide an approximate number of how many seeds each plant had produced, we generated two allometric relationships by visual counting of fruits per plant and seeds per fruit. In order to be sure that the counts corresponded to single plants, we counted fruits and seeds of only individual replicates of accessions, not the population replicates (see <u>Field experiment design section</u>). Because a strong relationship had already been validated between inflorescence size and the number of fruits in a number of studies with *A. thaliana*, we decided that counting a few inflorescences of three sizes, reflecting the broad size spectrum, would be sufficient to establish a first allometric relationship with the four image-acquired variables (n=11 inflorescences, R^2^=0.97, P=4×10^−4^, Fig.5B,C). To express fecundity as the number of seeds, we counted all seeds inside one fruit for each of the inflorescences used for the first allometric relationship (n=11 fruits), aiming for a wide range of fruit sizes. The mean was 28.3 seeds per fruit and the standard deviation was 11.2 seeds. The two aforementioned allometric relationships were used to predict, first, the number of fruits per inflorescence using the four image analysis variables, and second, the number of seeds corresponding to the number of fruits per inflorescence.

## Data records

The main datasets of accession information and trait values measured in the field for all replicates as well as curated averages per genotype are available as supplementary information at xxx. The datasets are also part of the R package “dryAR” available at http://github.com/MoisesExpositoAlonso/dryAR with doi xxx.

**Table 2.**
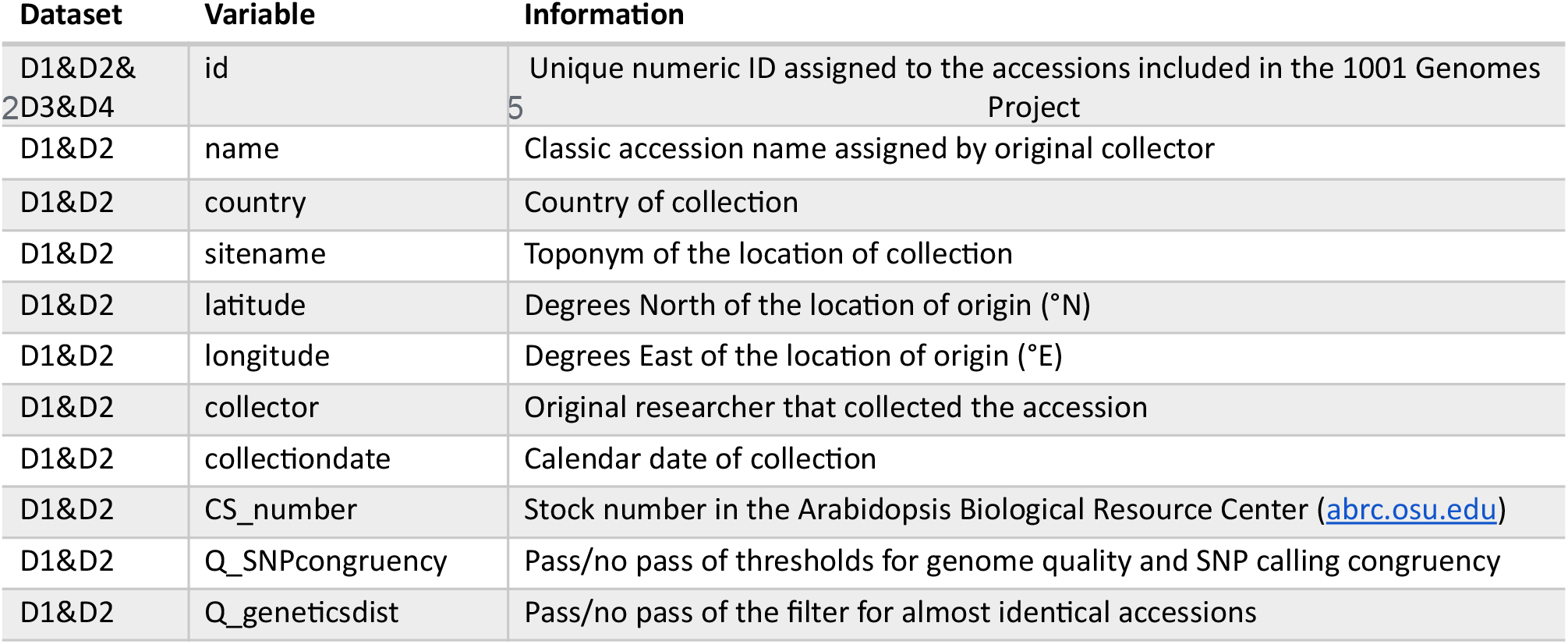
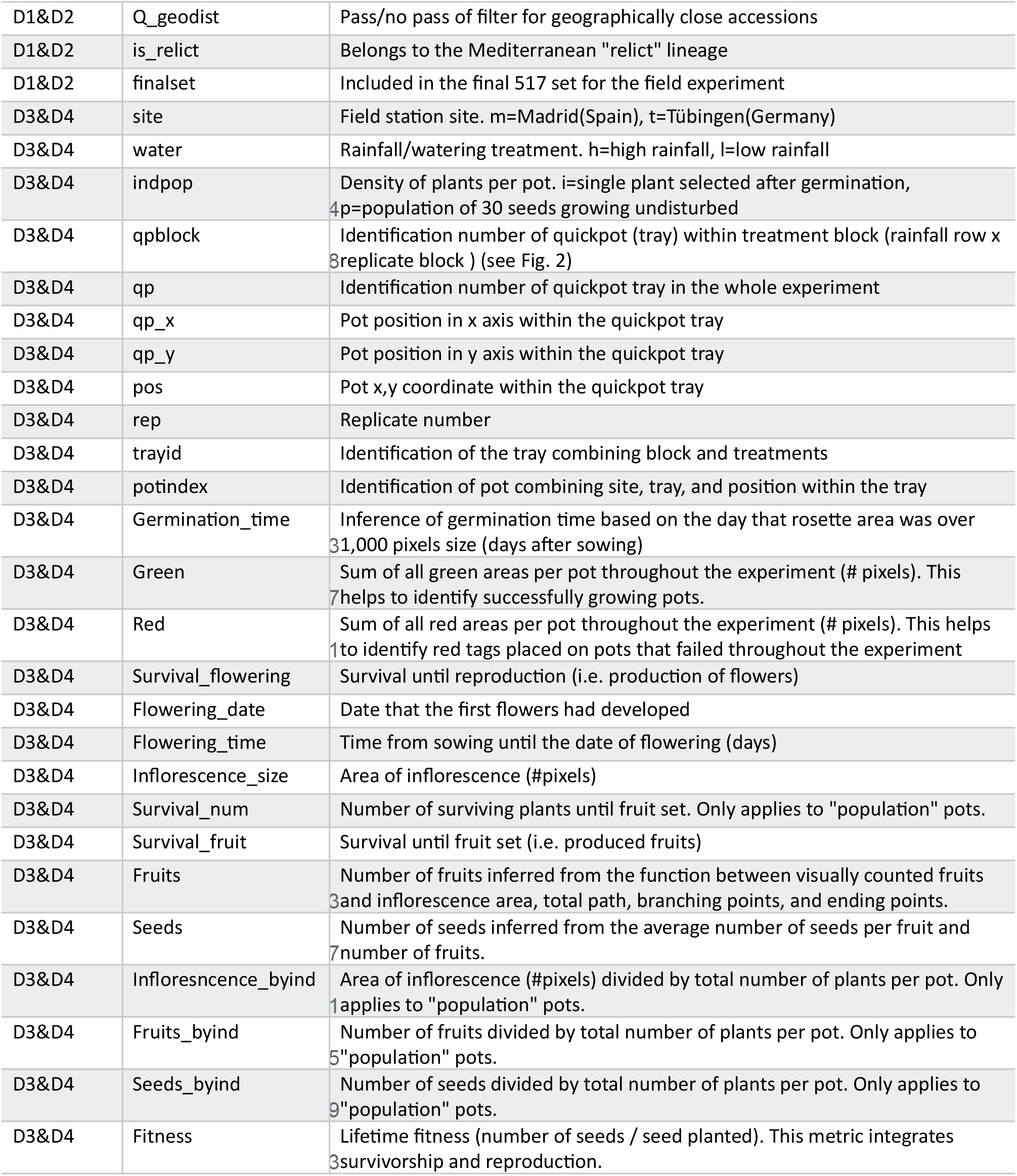
Variable descriptions. Variable names and their descriptions and units are reported. All datasets share a common accession identification number.

## Technical validation

### Data processing

All images are deposited at [updatehere]. The Python modules to process images for green area segmentation and inflorescence analyses are available at http://github.com/MoisesExpositoAlonso/hippo and http://github.com/MoisesExpositoAlonso/hitfruit, along with example datasets.

To reproduce our data curation procedure we created the R package dryAR (http://github.com/MoisesExpositoAlonso/dryAR with doi xxx). All scripts to re-generate data from raw files can be found at http://github.com/MoisesExpositoAlonso/dryAR/data-cleaning.

### Replicability of image processing

After testing different camera parameters, we used an exposure of −2/3 and an ISO of 100. White balance was set for flashlight. We used a dark box with all sides closed, so the flashlight was the only source of illumination. This ensured that the white balance and illumination were virtually consistent from picture to picture, as shown before^20^. Photos were saved both in .jpeg and .raw to allow for *a posteriori* adjustments if needed. Using a calibration board with 1.3 cm × 1.3 cm white and dark squares, we examined the error between the inferred area from image analysis and the real 1.3 cm-side squares across the tray. This provided us with a median resolution estimate of 101.5 pixels mm. The deviations from the true area were minimal, with a median of 2.7% and values of 1.4% / 4.2% for the 1^st^ and 3^rd^ quartile. The maximum area deviations were of 8 to 9% in the extreme corners of the tray, where we did not sow any seeds. We are confident that such small variation in retrieved area is compensated by the randomized locations of genotypes within the trays.

To further verify that our camera settings and segmentation pipeline produced replicable extractions of plant green area, we used images of trays that were photographed twice on the same day by mistake. In total there were 1,508 such pots distributed across 11 timepoints and different trays. By comparing the area of the same pot of two different camera shots and segmentation analyses, we could verify that the Spearman’s rank correlation was very high (r=0.97, n=1508, *P*<10^−16^), confirming high replicability.

Because we ran the same segmentation and skeletonization software on both rosette and inflorescence images, we could leverage the clearly different image patterns that rosettes and inflorescences have to identify labeling errors (i.e. mistakes in manually inpuffing sample information of the pictures). To do this, we first trained a random forest model to predict the manually labeled “rosette” or “inflorescence” by the four image variables in Fig. 5. By fitting a Random Forest with all images, we find that the leave-one-out accuracy was 92.1%, i.e. ca. 2,000 images were incorrectly labeled by the algorithm. We manually checked whether these were mislabeled or rather whether they “looked similar” in terms of area or landmark points in the photo, e.g. when both rosette or inflorescences were diminute. We found that only 2.5% were incorrectly mislabeled (and corrected them) and are thus confident that the labeling error must be below 2.5%.

### Experimental validation

Although repeating experiments in climatically-similar locations would be impractical, we could verify that survival in Madrid and low precipitation correlated with a preliminary drought experiment in the greenhouse (Spearman’s rho=0.17, n=211, *P*=0.01)^20^. On the other hand, reproductive allocation measured under optimal conditions in the greenhouse correlated with total seed output in the most similar field experiments, Tübingen high precipitatìon (Spearman’s rho=0.27, n=211, *P*=5×10^−5^)^23^.

## ADDITIONAL INFORMATION

### Acknowledgements

We are thankful to Belen Mendez-Vigo, Carlos Alonso-Blanco and the technical service at CBGP-UMP, Antolín López Quirós, Marisa López Herránz and Miguel Ángel Mora Plaza, for assistance during sowing in Madrid. We also thank Xavi Picó for advice on experimental design.

### Supplementary Information

accompanies this paper at xxx

### Author contributions

MEA conceived and designed the project. MEA carried out the experiment in Tübingen. MEA and RGR carried out the experiment in Madrid. All authors contributed to specific tasks in the experiments (see detailed description below). OB provided the field site in Tübingen and FGA provided the site in Madrid. DW secured funding for the project. MEA carried out the analyses and wrote the first draft of the manuscript. All authors edited, commented and approved the manuscript.

### Datasets

#### Dataset 1 Quality-based selection of the original 1,135 accessions

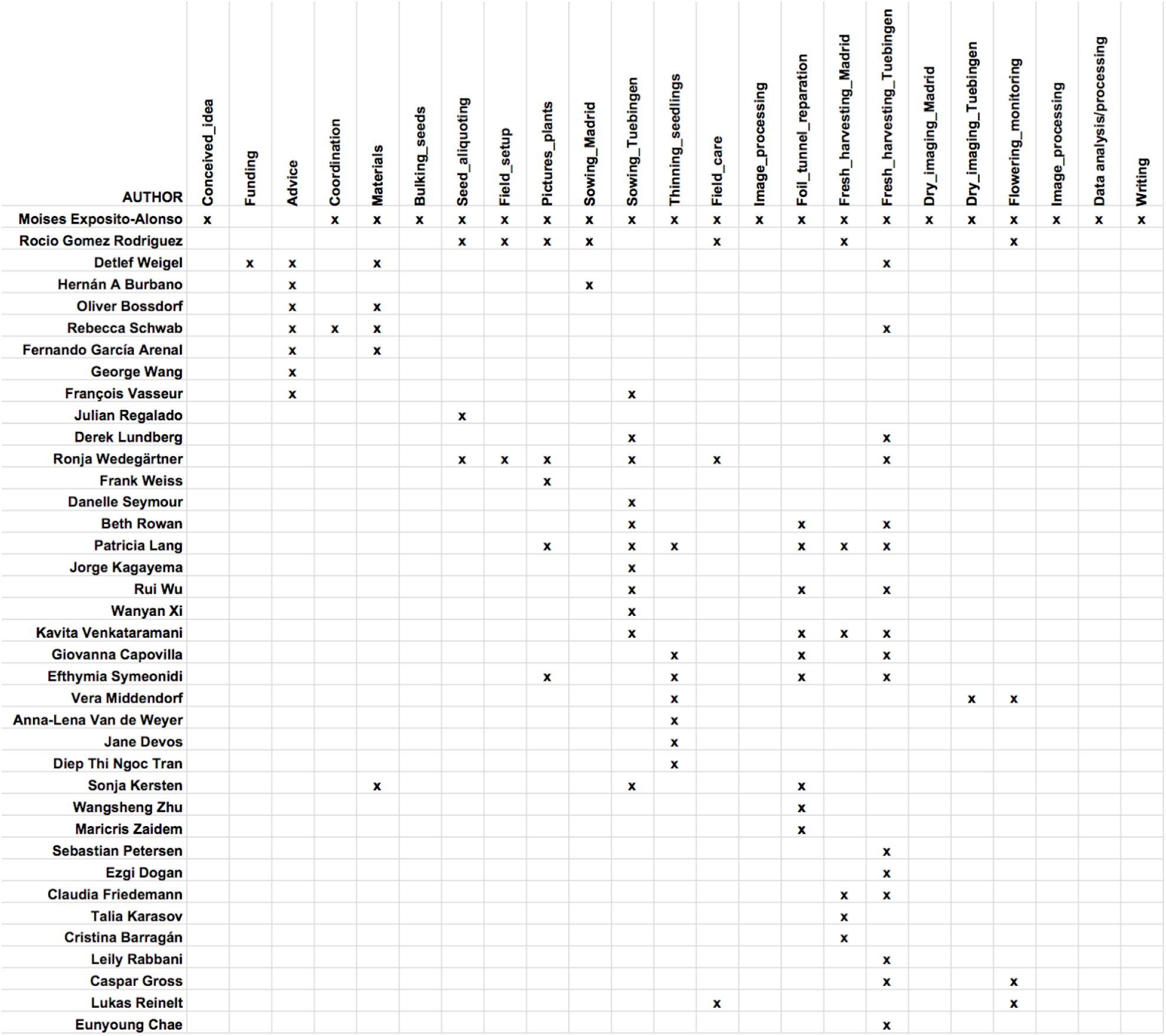

We report the 1001 Genome identification numbers, the quality filters that each accession passed during the selection of the 517 set.

#### Dataset 2 Description of the 517 accessions

We report the final set of 517 accessions that were used in the field experiment.

#### Dataset 3 All traits measured per replicate

For each pot replicate, we report all raw data as well as composite variables.

#### Dataset 4 Curated means per accession

For each accession, we report all data as well as composite variables.

